# Thymoquinone-Protoflavone Hybrid Molecules as Potential Antitumor Agents

**DOI:** 10.1101/2023.09.04.555971

**Authors:** Sara H. H. Ahmed, Bizhar A. Tayeb, Tímea Gonda, Gábor Girst, Kornél Szőri, Róbert Berkecz, István Zupkó, Renáta Minorics, Attila Hunyadi

**Affiliations:** Institute of Pharmacognosy, University of Szeged, H-6720 Szeged, Hungary; Department of Pharmacodynamics and Biopharmacy, University of Szeged, H-6720 Szeged, Hungary; Institute of Pharmaceutical Analysis, University of Szeged, 4, 6720 Szeged, Hungary; Interdisciplinary Centre of Natural Products, University of Szeged, H-6720 Szeged, Hungary

**Author notes:** **Corresponding Author** Attila Hunyadi. On leave from the Department of Pharmaceutical Chemistry, Faculty of Pharmacy, University of Khartoum, 11111 Khartoum, Sudan (Sara H. H. Ahmed).

## Abstract

We describe herein the synthesis of eight new ester-coupled hybrid compounds from thymoquinone and protoflavone building blocks, and their bioactivity testing against multiple cancer cell lines. Among the hybrids, compound 14 showed promising activities in all cell lines studied. The highest activities were recorded against breast cancer cell lines with higher selectivity to MDA-MB-231 as compared to MCF-7. Even though the hybrids were found to be completely hydrolysed in 24 h under cell culture conditions, compound 14 demonstrated a ca. three times stronger activity against U-87 glioblastoma cells than a 1:1 mixture of its fragments. Protoflavone-thymoquinone hybrids may therefore serve as potential new antitumor leads particularly against glioblastoma.

## Introduction

Cancer is the second leading cause of death worldwide [1]. Despite the many available therapeutic options including a wide range of chemotherapeutic agents of natural origin, many limitations exist for successful treatments, e.g., severe side effects and the frequent development of resistance [2]. Because of these, there is a continuous need for effective and safe new anticancer drugs. Among different types of cancer, Glioblastoma multiforme (GBM) is a very common and particularly aggressive tumour of the brain or the spinal cord, with a very poor prognosis and limited therapeutic options [3].

Protoapigenone (PA) is a fern-originated rare natural flavonoid that was found to be effective against many types of cancer *in vitro* and *in vivo* [4]. It inhibits the ataxia telangiectasia and Rad3 related (ATR)-dependent activation of checkpoint kinase 1 (Chk-1), which is a hallmark of DNA damage response (DDR) [5] and a promising antitumor target [6, 7] studied in many currently ongoing clinical trials, e.g., NCT02487095, NCT04616534, NCT04802174, NCT05338346, etc.

Thymoquinone (TQ), a monoterpene from the seeds of *Nigella sativa* L. (Ranunculaceae), was described as a promising antitumor lead based on its ability to regulate miRNAs expression influencing cell cycle progression, cell proliferation, metastasis, and angiogenesis [8, 9, 10, 11]. It also stimulated apoptotic genes in MDA-MB-231 cells by affecting cellular redox state [12]. Among other types of cancer, TQ was also studied against glioblastoma, and promising results were achieved. It promoted hydrogen peroxide generation, and disturbed cellular redox state and mitochondrial function leading to cell cycle arrest and apoptosis [13]. In addition, it inhibited autophagy and induced a caspase-independent glioblastoma cell death [14]. When combined with temozolomide (TMZ), a standard drug of glioblastoma treatment, TQ was reported to increase its efficacy [15, 16]. Interestingly, TQ-mediated apoptosis was found to occur due to a p53-mediated transcriptional repression of Chk-1 [17].

The design and preparation of hybrid molecules, i.e., the strategy of combining bioactive compounds into a single, supposedly multitarget entity, is an emerging concept in rational drug discovery [18]. This approach is especially attractive for complex diseases of a multifactorial profile like cancer [19, 20, 21, 22]. We have previously reported the preparation of two series of antitumor hybrids. Natural, or semi-synthetic protoflavonoids were linked to chalcone [23] or indole derivatives [24]. In both series, a protoflavone fragment was included as an ATR inhibitor, and it was combined with a fragment able to induce oxidative stress (i.e., a ferrocene or a chalchone), or to activate p53 (i.e., a spiropyrazole oxindole). Both hybrid compound series demonstrated greatly improved efficacy against breast cancer cell lines. In the current work our aim was to prepare new hybrid compounds of a protoflavone and TQ, i.e., to combine an ATR inhibitory fragment with one able to induce oxidative stress and interfere with p53-mediated Chk-1 activation, and to investigate their antiproliferative potentials against a cell line panel of gynecological or glioblastoma origin.

## Materials and Methods

### General Information

Reagents were purchased from Sigma (Merck KGaA, Darmstadt, Germany). Solvents (analytical grade for synthetic work and flash chromatography purifications and high-performance liquid chromatography (HPLC) grade for analytical and preparative HPLC work) were obtained from Chem-Lab NV (Zedelgem, Belgium), Macron Fine Chemicals (Avantor Performance Materials, Center Valley, PA, USA), VWR International S.A.S., and Fontenay-sous-Bois, France.

For purification, flash chromatography and/or RP-HPLC, was used. The former was performed on a CombiFlash Rf + Lumen apparatus (TELEDYNE Isco, Lincoln, NE, USA) equipped with evaporative light scattering (ELS) and diode array detectors. Teledyne Isco Inc. RediSep prefilled silica columns and cartidges were utilized. HPLC was conducted on an Armen Spot Prep II integrated HPLC purification system (Gilson, Middleton, WI, USA) with dual-wavelength detection, utilizing a Kinetex XB C18 (5 μm, 250 × 21.2 mm) column at a flow rate of 15 mL/min. Semi-preparative purification was performed on an Agilent 1100 series (Waters Co., Milford, MA, USA) connected to a Jasco UV-2075 detector (Jasco Co., Tokyo, Japan) utilizing a Gemini-NX C18 column (5 μm, 250 x 10 mm) and the flow rate was 3 mL/min. Solvent systems were selected for each compound based on their chromatographic behaviour on TLC.

The purity of the obtained compounds was assessed by RP-HPLC analyses on a system of two Jasco PU 2080 pumps, a Jasco AS-2055 Plus intelligent sampler connected to a JASCO LC-Net II/ADC equipped with a Jasco MD-2010 Plus PDA detector (Jasco International Co. Ltd., Hachioji, Tokyo, Japan) using a Kinetex C-18 (5 μm, 250 × 4.6 mm) column (Phenomenex Inc., Torrance, CA, USA) and applying a gradient of 30–100% aqueous AcN in 30 min followed by 100% AcN for another 10 min at a flow rate of 1 mL/min.

The ^1^H- and ^13^C-NMR spectra were measured in CDCl3, utilizing 5 mm tubes on a Bruker DRX-500 spectrometer at 500 (^1^H) and 125 (^13^C) MHz at room temperature with the deuterated solvent signal taken as reference. The standard Bruker pulse programs were used to the heteronuclear single quantum coherence (HSQC), heteronuclear multiple bond correlation (HMBC), ^1^H-^1^H correlation spectroscopy (COSY), and nuclear Overhauser effect spectroscopy (NOESY). High resolution mass spectroscopy (HRMS) was performed on a Q-Exactive Plus hybrid quadrupole-orbitrap mass spectrometer (Thermo Scientific, Waltham, MA, USA) equipped with a heated electrospray ionization (HESI-II) probe operated in positive or negative mode.

### Synthesis

#### Synthesis of 2-isopropyl-5-methylcyclohexa-2,5-diene-1,4-dione (2)

An aliquot of 2 g (0.013 mol) of compound **1** was dissolved in 90% aqueous AcN (50 mL) at room temperature, then (5.7 g, 0.013 mol) [Bis(trifluoroacetoxy)iodo]benzene (PIFA) was added. The reaction mixture was stirred for 1h, quenched, and solvents were evaporated under reduced pressure. The resulting mixture was directly purified using flash chromatography (Silica, gradient elution of 0–10% of EtOAc in n-hexane) to obtain compound **2** as a yellow crystalline solid (1.11 g, 50.6 %) [25].

#### General procedure for the synthesis of compounds 10-17

The method reported by Szakonyi et al. was used with modifications [26]. Compound **3** or **4** (50 mg, 0.21 mmol or 30 mg, 0.126 mmol) was dissolved in 2 ml of dry CH_2_Cl_2_ (5 mL), DMAP was added (2.6 mg, 0.021 mmol or 1.6 mg, 0.013 mmol), and the mixture was cooled to 0°C. Then, a solution of DCC in dry CH_2_Cl_2_ was added (44 mg, 0.21 mmol or 26 mg, 0.126 mmol). The mixture was stirred for 1 h at 0°C, after which the corresponding amount (1 eq.) of compound **6, 7, 8**, or **9** was added and left to stir overnight. The reaction mixture was washed with saturated NaHCO_3_ solution. The organic layer was collected, dried over Na_2_SO_4_, filtered, and the solvent was evaporated under reduced pressure. The resulting mixtures were purified using preparative RP-HPLC (Kinetex, C18, 5 μm, 250 × 21.2 mm). Some of them were subjected to further purification using semipreparative techniques (Agilent, C18, 5 μm, 250 × 10 mm) using appropriately selected aqueous AcN or MeOH solvent systems.

**Compound 3**. Yellow wax, 12.4%, C_13_H_16_O_4_, HRESIMS: [M+H]^+^ *m/z* = 237.11199, (calcd 237.11269); ^1^H NMR (500 MHz, in CDCl_3_): δ_H_ = 6.50 (s, 1H), 3.04 (hept, 1H, *J*=6.9 Hz), 2.83 (t, 2H, *J*=7.6 Hz), 2.52 (t, 2H, *J*=7.8 Hz), 2.06 (s, 3H), 1.11 (d, 6H, *J*=6.8 Hz) ppm; ^13^C NMR (125 MHz, fin CDCl_3_): δ_C_ = 188.17, 186.9, 177.5 (only detected on HSQC),154.9, 142.9, 141.5, 130.4, 32.6, 26.9, 22.4, 21.6, 11.9.

**Compound 4**. Yellow oil, 2.7%, C_13_H_16_O_4_, HRESIMS: [M+H]^+^ *m/z* = 237.11191, (calcd 237.11269); ^1^H NMR (500 MHz, in CDCl_3_): δ_H_ = 6.48 (s, 1H), 3.06 (hept, 1H, *J*=7.0 Hz), 2.85 (t, 2H, *J*=7.9 Hz), 2.48 (t, 2H, *J*=8.0 Hz), 2.01 (s, 3H), 1.28 (d, 6H, *J*=6.9 Hz) ppm; ^13^C NMR (125 MHz, in CDCl_3_): δ_C_ = 188.1, 187.7, 177.8, 149.9, 144.7, 141.9, 134.6, 33.4, 29.4, 21.6, 21.2, 15.7.

**Compound 10**. Yellow oil, 12.2 %, C_28_H_24_O_9_, HRESIMS: [M-H]^-^ *m/z* = 503.13453, (calcd 503.13421); ^1^H NMR (500 MHz, in CDCl_3_): δ_H_ = 12.39 (s, 1H), 6.85 (d, 2H, *J*=9.6 Hz), 6.72 (s, 1H), 6.66 (s, 1H), 6.56 (s, 1H), 6.52 (s, 1H), 6.41 (d, 2H, *J*=9.6 Hz), 3.05 (hept, 1H, *J*=7.0 Hz), 2.92 (t, 2H, *J*=7.7 Hz), 2.73 (t, 2H, *J*=7.7 Hz), 2.09 (s, 3H), 1.12 (d, 6H, *J*=6.8 Hz) ppm; ^13^C NMR (125 MHz, in CDCl_3_): δ_C_ = 188.0, 186.9, 184.4, 182.9, 170.1, 166.4, 162.1, 156.8, 156.2, 154.9, 2 × 145.3, 142.4, 141.7, 2 × 130.6, 130.5, 109.0, 107.9, 106.0, 101.3, 69.7, 33.0, 26.9, 22.4, 21.6, 12.1.

**Compound 11**. Yellow oil, 8.3 %, C_29_H_26_O_9,_ HRESIMS: [M+H]^+^ *m/z* = 519.16564, (calcd 519.16551); ^1^H NMR in (500 MHz, in CDCl_3_): δ_H_ = 12.43 (s, 1H), 6.75 (d, 2H, *J*=9.7 Hz), 6.70 (s, 1H), 6.63 (s, 1H), 6.58 (s, 1H), 6.56 (s, 1H), 6.53 (d, 2H, *J*=10.9 Hz), 3.41 (s, 3H), 3.05 (hept, 1H, *J*=6.9 Hz), 2.92 (t, 2H, *J*=7.7 Hz), 2.72 (t, 2H, *J*=7.7 Hz), 2.09 (s, 3H), 1.12 (d, 6H, *J*=6.8 Hz) ppm; ^13^C NMR (125 MHz, in CDCl_3_): δ_C_ = 188.0, 186.9, 184.3, 182.9, 170.1, 165.4, 162.1, 156.7, 156.2, 154.9, 2 × 145.1, 142.4, 141.7, 2 × 133.6, 130.5, 109.1, 108.5, 105.9, 101.2, 74.9, 52.9, 33.0, 26.9, 22.4, 21.6, 12.1.

**Compound 12**. Yellow oil, 8.0 %, C_30_H_28_O_9_, HRESIMS: [M+H]^+^ *m/z* = 533.18104, (calcd 533.18116); ^1^H NMR (500 MHz, in CDCl_3_): δ_H_ = 12.44 (s, 1H), 6.79 – 6.74 (m, 3H), 6.62 (s, 1H), 6.56 – 6.51 (m, 4H), 3.58 (q, 2H, *J*=6.9 Hz), 3.05 (hept, 1H, *J*=6.9 Hz), 2.92 (t, 2H, *J*=7.7 Hz), 2.72 (t, 2H, *J*=7.7 Hz), 2.09 (s, 3H), 1.29 (t, 3H, *J*=6.9 Hz), 1.12 (d, 6H, *J*=6.8 Hz) ppm; ^13^C NMR (125 MHz, in CDCl_3_): δ_C_ = 188.0, 186.9, 184.5, 182.9, 170.1, 165.7, 162.1, 156.7, 156.1, 154.9, 2 × 145.6, 142.4, 141.7, 2 × 133.1, 130.5, 109.1, 108.6, 105.9, 101.2, 74.6, 61.1, 33.0, 26.9, 22.4, 21.6, 15.8, 12.1.

**Compound 13**. Yellow oil, 13.6 %, C_32_H_32_O_9_, MS: [M+H]^+^ *m/z* = 561.4, (calcd 561.2); ^1^H NMR (500 MHz, in CDCl_3_): δ_H_ = 12.45 (s, 1H), 6.80 – 6.72 (m, 3H), 6.62 (s, 1H), 6.57 – 6.50 (m, 4H), 3.51 (t, 2H, *J*=6.3 Hz), 3.05 (hept, 1H, *J*=7.0 Hz), 2.92 (t, 2H, *J*=7.7 Hz), 2.72 (t, 2H, *J*=7.3 Hz), 2.09 (s, 3H), 1.68 – 1.58 (m, 2H), 1.48 – 1.39 (m, 2H), 1.12 (d, 6H, *J*=6.9 Hz), 0.94 (t, 3H, *J*=7.3 Hz) ppm; ^13^C NMR (125 MHz, in CDCl_3_): δ_C_ = 188.0, 186.9, 184.5, 182.9, 170.1, 165.7, 162.1, 156.7, 156.1, 154.9, 2 × 145.7, 142.4, 141.7, 2 × 133.1, 130.5, 109.1, 108.6, 105.9, 101.2, 74.6, 65.2, 33.0, 32.2, 26.9, 22.4, 21.6, 19.4, 13.9, 12.1.

**Compound 14**. Yellow oil, 10.9 %, C_28_H_24_O_9_, HRESIMS: [M+H]^+^ *m/z* = 505.15037, (calcd 505.14986); ^1^H NMR (500 MHz, in CDCl_3_): δ_H_ = 12.37 (s, 1H), 6.86 (d, 2H, *J*=10.1 Hz), 6.72 (s, 1H), 6.67 (d, 1H, *J*=2.0 Hz), 6.58 (d, 1H, *J*=2.0 Hz), 6.50 (d, 1H, *J*=1.6 Hz), 6.41 (d, 2H, *J*=10.0 Hz), 3.08 (hept, 1H, *J*=7.0 Hz), 2.95 (t, 2H, *J*=7.1 Hz), 2.68 (t, 2H, *J*=7.1 Hz), 2.02 (d, 3H, *J*=1.6 Hz), 1.31 (d, 6H, *J*=7.0 Hz) ppm; ^13^C NMR (125 MHz, in CDCl_3_): δ_C_ = 188.2, 187.5, 184.3, 182.9, 169.9, 166.3, 162.1, 156.8, 156.4, 150.1, 2 × 145.2, 144.7, 141.5, 134.7, 2 × 130.7, 109.1, 108.0, 106.1, 101.2, 69.8, 33.9, 29.5, 21.7, 21.3, 15.7.

**Compound 15**. Yellow oil, 25.8 %, C_29_H_26_O_9,_ HRESIMS: [M+H]^+^ *m/z* = 519.16604, (calcd 519.16551); ^1^H NMR (500 MHz, in CDCl_3_): δ_H_ = 12.41 (s, 1H), 6.75 (d, 2H, *J*=10.3 Hz), 6.70 (s, 1H), 6.64 (d, 1H, *J*=2.1 Hz), 6.58 – 6.54 (m, 3H), 6.50 (d, 1H, *J*=1.6 Hz), 3.41 (s, 3H), 3.08 (hept, 1H, *J*=7.0 Hz), 2.95 (t, 2H, *J*=7.1 Hz), 2.67 (t, 2H, *J*=7.1 Hz), 2.02 (d, 3H, *J*=1.5 Hz), 1.31 (d, 6H, *J*=7.0 Hz) ppm; ^13^C NMR (125 MHz, in CDCl_3_): δ_C_ = 188.1, 187.5, 184.2, 182.9, 169.8, 165.5, 162.1, 156.8, 156.3, 150.1, 2 × 144.9, 144.7, 141.5, 134.7, 2 × 133.6, 109.2, 108.6, 105.9, 101.2, 75.1, 52.9, 33.9, 29.5, 21.7, 21.3, 15.7.

**Compound 16**. Yellow oil, 13.2%, C_30_H_28_O_9_, HRESIMS: [M+H]^+^ *m/z* = 533.18172, (calcd 533.18116); ^1^H NMR (500 MHz, in CDCl_3_,): δ_H_ = 12.42 (s, 1H), 6.77 (d, 2H, *J*=10.3 Hz), 6.75 (s, 1H), 6.63 (d, 1H, *J*=2.0 Hz), 6.56 (d, 1H, *J*=2.1 Hz), 6.53 (d, 2H, *J*=10.2 Hz), 6.50 (d, 1H, *J*=1.6 Hz), 3.59 (q, 2H, *J*=7.0 Hz), 3.08 (hept, 1H, *J*=7.0 Hz), 2.95 (t, 2H, *J*=7.1 Hz), 2.67 (t, 2H, *J*=7.1 Hz), 2.02 (d, 3H, *J*=1.6 Hz), 1.32 – 1.2 (m, 9H) ppm; ^13^C NMR (125 MHz, in CDCl_3_): δ_C_ = 188.1, 187.5, 184.4, 182.9, 169.9, 165.7, 162.1, 156.8, 156.2, 150.1, 2 × 145.5, 144.7, 141.5, 134.7, 2 × 133.1, 109.2, 108.6, 105.9, 101.2, 74.7, 61.2, 33.9, 29.4, 21.7, 21.3, 15.8, 15.7.

**Compound 17**. Yellow oil, 11.3%, C_32_H_32_O_9_, HRESIMS: [M+H]^+^ *m/z* = 561.21316, (calcd 561.21246); ^1^H NMR (in CDCl_3_, 500 MHz): δ_H_ = 12.45 (s, 1H), 6.77 – 6.73 (m, 3H), 6.63 (d, 1H, *J*=2.0 Hz), 6.56 (d, 1H, *J*=2.1 Hz), 6.53 (d, 2H, *J*=10.1 Hz), 6.50 (d, 1H, *J*=1.6 Hz), 3.52 (t, 2H, *J*=6.3 Hz), 3.07 (hept, 1H, *J*=7.0 Hz), 2.94 (t, 2H, *J*=8.8 Hz), 2.67 (t, 2H, *J*=8.8 Hz), 2.02 (d, 3H, *J*=1.6), 1.67 – 1.59 (m, 2H), 1.48 – 1.39 (m, 2H), 1.31 (d, 6H, *J*=7.0 Hz), 0.95 (t, 3H, *J*=7.4 Hz) ppm; ^13^C NMR (in CDCl_3_, 125 MHz): δ_C_ = 188.1, 187.6, 184.5, 183.0, 169.9, 165.7, 162.1, 156.7, 156.2, 150.1, 2 × 145.7, 144.7, 141.5, 134.6, 2 × 133.1, 109.1, 108.6, 105.9, 101.3, 74.6, 65.2, 33.8, 32.2, 29.5, 21.6, 21.3, 19.4, 15.7, 13.9.

#### Enzymatic Hydrolysis assay

A 0.1 M solution of compound **11** or **15** in AcN was prepared and added to 0.025 M PBS (pH=7.4) equilibrated in a water bath at 37°C. (170 units/ mg protein). Porcine esterase (lypholized powder, Sigma Aldrich, St. Louis, Co., USA) was diluted with 0.025 M PBS then this volume was completed to 2.5 mL with PBS to result in a final compound concentration of 8×10^−4^ M and 1.3 units of enzyme/ml. After 24 hrs incubation at 37 °C, the enzyme activity was quenched and the samples were analysed via RP-HPLC (Kinetex, C18, 5 μm, 250 × 4.5 mm column, 30-100% AcN gradient elution) [27].

#### Cell lines and culture conditions

Gynecological cancer cell lines of human origin including breast cancer like the triple negative MDA-MB-231 and estrogen receptor positive MCF-7, HPV16-positive cervical adenocarcinoma (HeLa), and human glioblastoma (U-87) cell lines were used as *in vitro* models to study the antiproliferative effects of the evaluated compounds. All cell lines were cultivated in T-75 flasks in a minimal essential medium (MEM) supplemented with 10% heat-inactivated fetal bovine serum (FBS), 1% antibiotic-antimycotic mixture (penicillin-streptomycin-amphotericin B) and 1% non-essential amino acids. The cells were incubated at 37°C in 5% CO^2^ incubator. The cells were seeded in 96-well plates at a density of 5 × 10^3^ in 100 μL per well, except for the U-87 that were seeded in 1 × 10^4^ and incubated at the same conditions for overnight to allow the cells’ attachment to the well’s bottom before the treatment.

#### *In vitro* antiproliferative assay

The compounds were dissolved in dimethyl sulfoxide (DMSO) as 10 mM standard stock solutions and kept at -20°C with minimum light exposure. Immediately before each experiment, the stock solution was used and diluted with a culture medium to get the final concentrations. The values of half-maximal inhibitory concentration (IC_50_) were determined by exposure of the cells into eight different concentrations of each tested compound (0.39, 0.78, 1.56, 3.125, 6.25, 12.5, 25 and 50 μM). Temozolomide (TMZ) and cisplatin were used as positive controls in the case of U-87 and the gynecological cell lines, respectively. The negative control wells included the cells with only MEM treatment. The plates were incubated for up to 72 hours under the same abovementioned incubation conditions. The colorimetric MTT assay was used to assess the compounds’ effect on cell proliferation. Briefly, 20 μL of MTT solution ([3-(4,5-dimethylthiazol-2-yl)-2,5-diphenyltetrazolium bromide], 5 mg/mL in PBS, Duchefa Biochemie BV, Haarlem, The Netherlands) was added to each well including the negative controls and kept under the usual incubation circumstances for additional four hours. Subsequently, the media was carefully aspirated and 100 μL of DMSO was added to each well and the plates were gently shaken for 30 min to solubilize the precipitated crystals of purple formazan. Absorbance was measured at a wavelength of 545 nm using a microplate UV-VIS reader (SPECTROstar Nano, BMG Labtech GmbH, Offenburg, Germany).(28)

#### Combination Assay

Combination study was performed by treating cells with equimolar mixtures of the abovementioned fragments or their hybrids, and corresponding cell viability data were comparatively evaluated. In this bioassay, at least two separate experiments were performed, each in triplicate. The dataset was then subjected to appropriate statistical analysis. The calculated IC_50_ values were subsequently employed to quantitatively assess the extent of pharmacological benefit obtained through the hybridization of fragments in comparison to the cytotoxic effect produced by the experimental combination of PA or its derivatives and TQ fragments.

#### Nonlinear regression and statistical analysis

Cell viability data were collected from two separate experiments in triplicates and evaluated using GraphPad Prism 9.5.1 (GraphPad Software Inc., San Diego, CA, USA). The half-maximal inhibitory concentration (IC_50_) values were determined using the log inhibitor vs normalized response nonlinear regression model. Difference between the IC_50_ values of a hybrid and its corresponding fragments’ experimental combination was statistically evaluated using unpaired T-test.

## Results and Discussion

### Chemistry

In this work, TQ (**2**) was synthesized from thymol (**1**) using two different methods. In our case, the first procedure described by Asakawa et al. [29] required multiple purification steps due to the co-elution of thymol (**1**) with TQ. This negatively affected the yield (24.2%). Attempts to optimize the purification were not successful, however, the use of polyamide as a stationary phase and DCM as a mobile phase in a second purification step was, to some extent, helpful.

Using our method previously used for the preparation of protoflavonoids [25] resulted in a better yield (up to 50.6%) with a single purification step, and, to our knowledge, this is the first report for the synthesis of TQ from thymol using PIFA as an oxidizing agent.

**Scheme 1.**
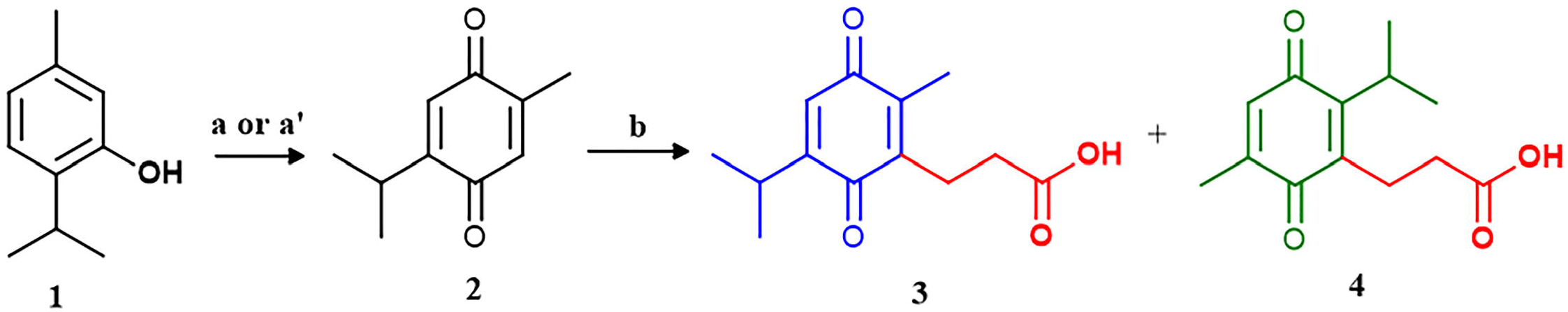
Preparation of TQ (2) and its derivatives; compounds 3 and 4. Reaction conditions: a. mCPBA/CHCl_3_/rt; a’ . PIFA/AcN: H_2_O; 9:1/rt/1h; b. succinic acid/ AgNO_3_/(NH_4_)_2_S_2_O_8_/AcN/H_2_O/100°C.

To link TQ and PA; reacting TQ with succinic acid resulted in compounds **3** and **4** as an isomeric mixture, and the isomers were isolated by preparative RP-HPLC using a Kinetex XB C18 (5 μm, 250 × 21.2 mm) column and 50% aqueous MeOH containing 0.1% formic acid. To our knowledge, this is the first time to report that an isomeric mixture resulted from this reaction and to successfully separate the isomers. Compounds **3** and **4** were then esterified with different protoflavones to yield eight compounds (4 isomeric pairs). Each pair is different from the others by the substituent at the 1□-position of the protoflavone’s B-ring (**Scheme 2**).

**Scheme 2.**
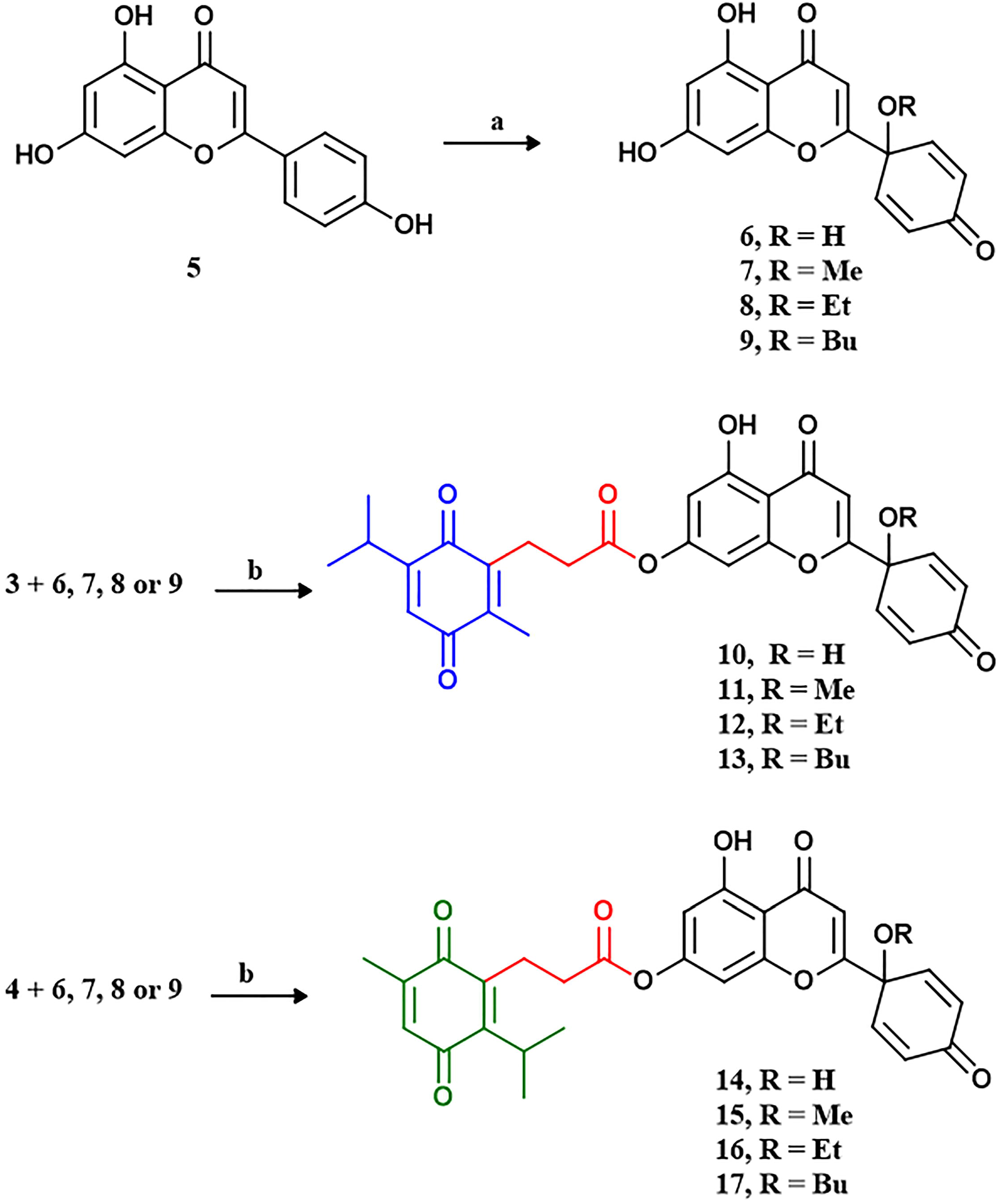
Preparation of the protoflavone fragments (6-9) and the hybrids (10-17). Reaction conditions: a. PIFA/AcN: ROH; 9:1/70oC/1h; b. DDC/DMAP/dry DCM/0 °C.

The structure of all target compounds was confirmed by HRMS and 1D and 2D NMR spectroscopy. All protoflavones as well as the hybrids possessed the aromatic region peaks characteristic of protoflavone B-ring hydrogens with a ^1^H-^1^H coupling constant of around 10 Hz. In the case of compound **3** the carboxylic quaternary C peak could only be detected in the HSQC spectra.

### *In vitro* antiproliferative assay

The antiproliferative activity of PA (**6**) was found to be superior to that of TQ (**2**) across all tested cell lines (**Table 1**). Additionally, all hybrid compounds demonstrated potent antiproliferative effects against the cell lines assessed, typically with IC_50_ values lower than 10 μM, with the sole exception of compound **15** on U-87 cells. Moreover, their cell proliferation-inhibiting effects were found comparable or stronger than that of the utilized positive control compounds, cisplatin and temozolomide. Among the hybrid compounds, **14** exhibited the highest efficacy against cancer cells, with IC_50_ values ranging from 0.51 to 1.20 μM. In MDA-MB-231 cells, this activity seems to be primarily due to the protoflavone fragment (**6**) included in compound **14**, based on the identical IC_50_ values for these two compounds. In case of the other three cell lines, however, **14** behaved differently than **6**, This indicates that the hybrid coupling with TQ modified the protoflavone’s cell line specificity: HeLa and U-87 cells were more sensitive to the hybrid than to the protoflavone alone. Intriguingly, compound **14** showed selectivity towards MDA-MB-231 vs. MCF-7 cells, even though this was rather due to the higher resistance of the latter (**Fig 1**). Nevertheless, this demonstrates the appearance of a characteristic property of TQ in the hybrid compound’s pharmacological behavior, i.e., TNBC selectivity (**Table 1**).

**Table 1.** Antiproliferative effects of thymoquinone-protoflavone hybrids (10-17) and their building blocks (2-4 and 6-9) on cancer cell lines.

|  | Calculated IC<sub>50</sub> ± SEM; [μM]<sup>a</sup> |  |  |  |
| --- | --- | --- | --- | --- |
| Compound | MDA-MB-231 | MCF-7 | HeLa | U-87 |
| **2** | 7.02 ± 0.17 | 23.97 ± 1.37 | > 100 | 39.07 ± 3.53 |
| **3** | 38.17 ± 2.81 | > 100 | > 100 | 77.72 ± 1.61 |
| **4** | 12.44 ± 0.37 | > 100 | > 100 | 88.73 ± 5.17 |
| **6** | 0.57 ± 0.07 | 0.66 ± 0.06 | 1.80 ± 0.09 | 1.73 ± 0.11 |
| **7** | 2.23 ± 0.13 | 3.95 ± 0.30 | 5.51 ± 0.21 | 7.03 ± 0.11 |
| **8** | 1.22 ± 0.03 | 2.50 ± 0.11 | 2.83 ± 0.12 | 1.73 ± 0.06 |
| **9** | 0.82 ± 0.05 | 2.01 ± 0.07 | 1.88 ± 0.10 | 1.50 ± 0.12 |
| **10** | 1.27 ± 0.04 | 1.65 ± 0.07 | 2.00 ± 0.26 | 6.16 ± 0.49 |
| **11** | 2.25 ± 0.13 | 2.66 ± 0.11 | 3.51 ± 0.17 | 7.63 ± 0.46 |
| **12** | 2.15 ± 0.08 | 3.21 ± 0.07 | 2.35 ± 0.15 | 8.22 ± 0.73 |
| **13** | 0.99 ± 0.05 | 1.68 ± 0.16 | 1.40 ± 0.25 | 3.43 ± 0.25 |
| **14** | 0.52 ± 0.02 | 1.20 ± 0.03 | 1.06 ± 0.08 | 1.16 ± 0.20 |
| **15** | 3.53 ± 0.17 | 5.44 ± 1.32 | 6.78 ± 0.28 | 18.89 ± 3.42 |
| **16** | 1.98 ± 0.06 | 4.11 ± 0.34 | 1.71 ± 0.16 | 6.06 ± 0.40 |
| **17** | 1.075 ± 0.08 | 2.70 ± 0.09 | 3.08 ± 0.46 | 8.65 ± 1.15 |
| **Cis<sup>b</sup>** | 9.71 ± 0.51 | 6.55 ± 0.77 | 16.01 ± 2.00 | 9.13 ± 1.79 |
| **TMZ<sup>b</sup>** | - | - | - | 388.2 ± 43.0 |

<sup>a</sup> Mean value from two independent measurements with three replicates each
<sup>b</sup> Positive control; Cis: cisplatin, TMZ: temozolomide

**Figure 1.**
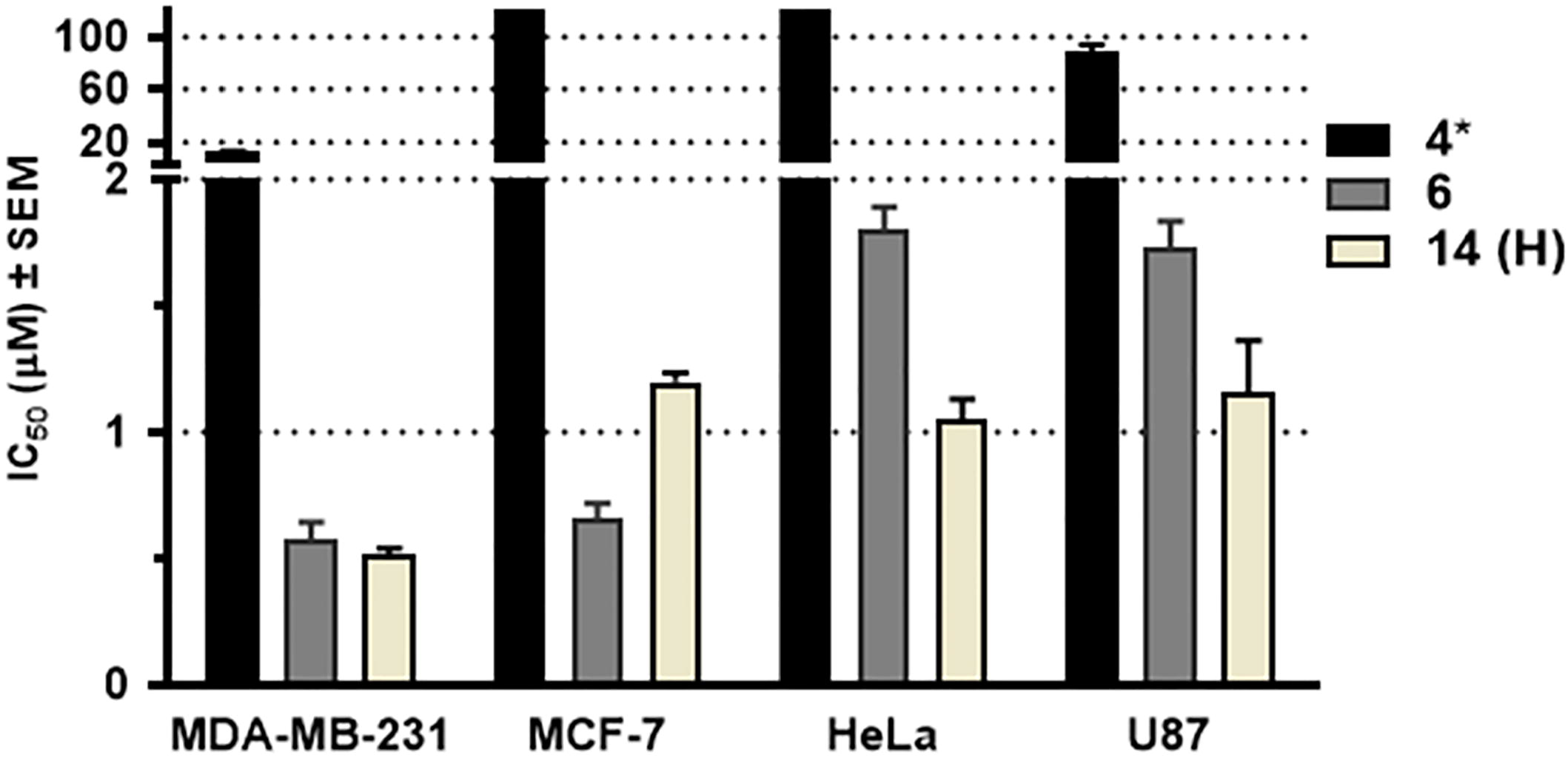
Calculated IC_50_ values of the hybrid; compound 14 and its building blocks (4 and 6) on the tested cancer cell lines. *Calculated IC_50_ values of compound 4 on MCF-7 and HeLa cells were above 100 μM.

Concerning structure-activity relationships (SAR) of these compounds, our results come in accordance with our previous findings with respect to the pattern of the protoflavones activity on breast cancer cell lines [25]. Accordingly, PA was found more potent on MCF-7 and MDA-MB-231 cells than its methylated derivative (**7**), and the activity was restored when increasing the length of the 1′-O-alkyl side-chain. The SAR of protoflavones followed a similar pattern on HeLa and U-87 cell lines. To some extent, this pharmacological behaviour could also be observed in case of the hybrids, even though the most potent compound was undoubtedly the 1′-OH substituted compound **14**.

The effect of isomerism on the activity seems a more complicated case. It appears that hybrids follow different SAR depending on the 1′ substituent of the protoflavone fragment. Accordingly, 1′-O-alkyl compounds **11**–**13** are generally more potent than their respective isomeric pairs **15**–**17**, while in case of the 1′-OH substitution the other isomer, **14** is the preferable one over its pair **10**.

Considering the hydrolysable nature of the ester coupling, it was of interest to test the stability of the compounds in the presence of esterase enzyme, as well as under cell culture conditions. After a 24h treatment with porcine liver esterase, a complete hydrolysis of the hybrids was observed. Further, the same was observed in MEM medium without enzymatic treatment. This suggests that the hybrids’ chemical stability needs to be improved for a possible further development.

Our next step was to evaluate if the hybrids merely act as pro-drugs of their fragments or if the fragments’ coupling into hybrids has a relevant pharmacodynamic benefit. To this, antiproliferative activity of a total of eight combinations (i.e., 1:1 mixture of building blocks) were tested on the U-87 cells in comparison with the corresponding hybrids. Results are compiled in **Table 2**.

**Table 2.**
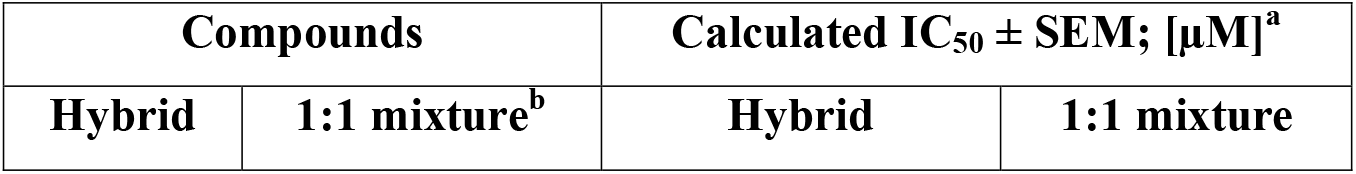
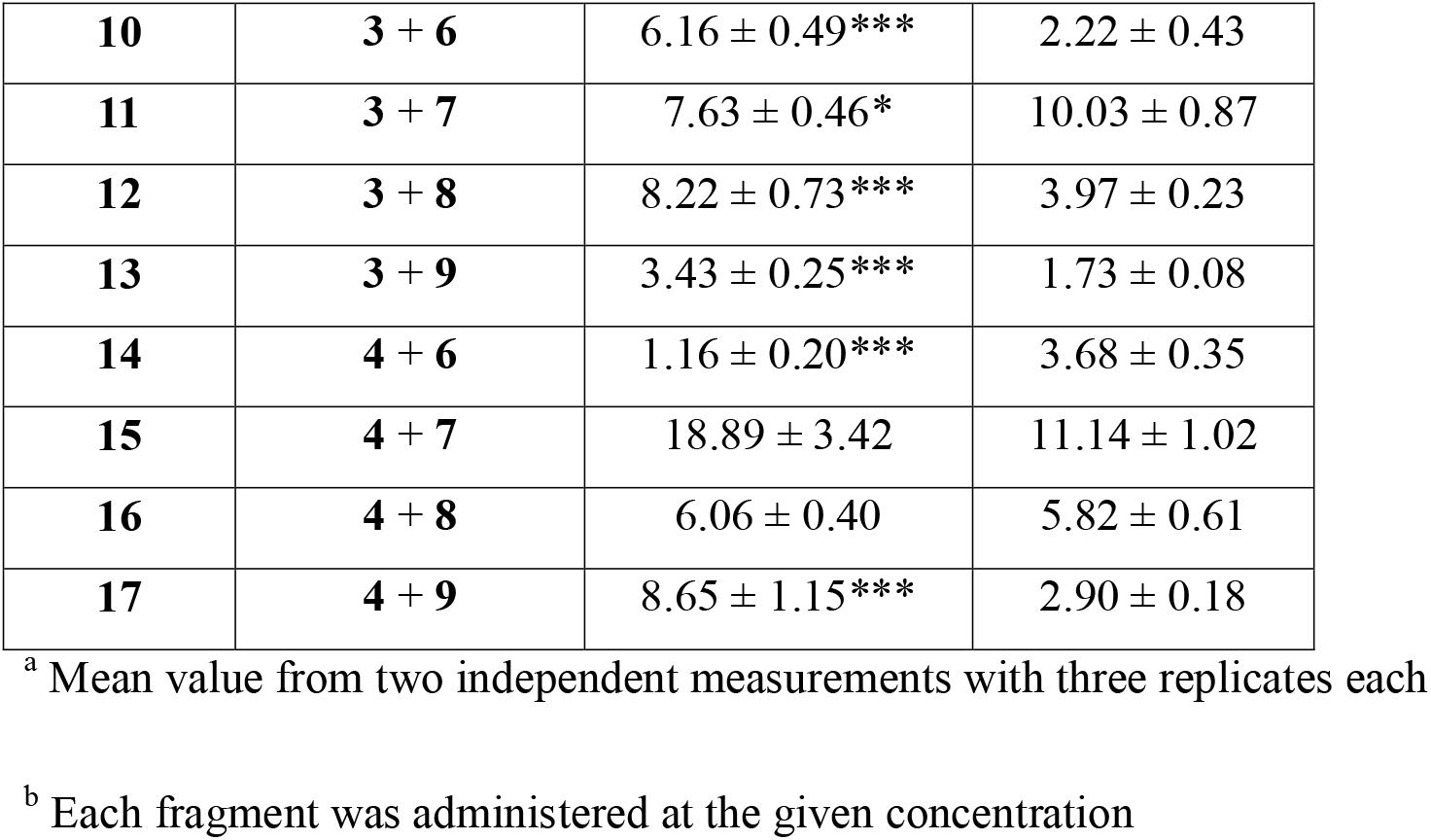
Calculated IC_50_ values of the structural combination (hybrid compounds) and experimental combination of the corresponding thymoquinone and protoflavone building blocks on the U-87 cells. Statistical analysis was performed by using unpaired t-test, * p<0.05; *** p<0.001 as compared to the corresponding 1:1 fragment mixture.

The term “synergism” in the context of a structural combination of fragments, i.e., a hybrid compound, indicates that it exhibits a significantly enhanced antiproliferative activity that surpasses the sum of the individual effects exhibited by its fragments. This was the case for two promising hybrids (**14** and **11**) and their building blocks, **4**:**6** and **3**:**7**, respectively. In particular, the most potent hybrid compound **14** exhibited a significant three-fold stronger activity than the mixture of its fragments **4** and **6**. These findings support the synthesis of such hybrid compounds as a valid strategy against glioblastoma.

In four cases, the IC_50_ values of the experimental combinations showed at least two-fold difference when compared to that of the hybrid compounds. On the other hand, results observed for compounds **15** and **16** showed no apparent difference between the antiproliferative activity of the hybrids and that of the corresponding fragment mixtures.

By this time no further information is available on the reason behind the observed differences in the hybrids’ pharmacological behavior. Further studies are necessary to understand the mechanisms leading to the superior activity of compound **14**.

## Conclusions

The current study led to the identification of a potent antitumor thymoquinone-protoflavone hybrid (**14**). Cell line selectivity pattern of this compound indicates pharmacodynamic properties combining those of the fragments. The hydrolysable ester linker releases the fragments relatively fast, within one hour in cell culture medium. Nevertheless, compound **14** demonstrated a ca. three times higher efficacy against U-87 glioblastoma cells than a co-treatment with the fragments. This strongly suggests that hybrid compounds of these two fragments may serve as potential new leads against glioblastoma, and the synthesis of more stable analogues, and/or development of appropriate formulations improving the stability of compound **14** is warranted.

## Author information

## Author Contributions

The manuscript was written through contributions of all authors. All authors have given approval to the final version of the manuscript.

## Acknowledgment

The authors would like to acknowledge Márton Benedek Háznagy and Gordana Krstic for their appreciated help in the enzymatic assay and NMR investigations, respectively.

## Abbreviations

TLC: thin layer chromatography

## Supporting Information

HRMS (**S1**-**S10**), ^1^H and ^13^C-NMR (**S11**-**S30**) spectra for compounds **3**,**4** and **10**-**17** are available.

