## Supporting Information for "Thymoquinone-Protoflavone Hybrid Molecules as Potential Antitumor Agents"

**Table of contents**

Figure S1. HRMS spectrum of compound 3

Figure S2. HRMS spectrum of compound 4

Figure S3. HRMS spectrum of compound 10

Figure S4. HRMS spectrum of compound **11**

Figure S5. HRMS spectrum of compound 12
Figure S6. HRMS spectrum of compound 13
Figure S7. HRMS spectrum of compound 14
Figure S8. HRMS spectrum of compound 15
Figure S9. HRMS spectrum of compound 16

Figure S10. HRMS spectrum of compound 17

Figure S11. ^1^H-NMR spectrum of compound 3

Figure S12. ^13^C-NMR spectrum of compound **3**

Figure S13. ^1^H-NMR spectrum of compound 4
Figure S14. ^13^C-NMR spectrum of compound 4
Figure S15. ^1^H-NMR spectrum of compound 10
Figure S16. ^13^C-NMR spectrum of compound 10
Figure S17. ^1^H-NMR spectrum of compound 11
Figure S18. ^13^C-NMR spectrum of compound 11
Figure S19. ^1^H-NMR spectrum of compound 12
Figure S20. ^13^C-NMR spectrum of compound 12

Figure S21. ^1^H-NMR spectrum of compound 13

Figure S22. ^13^C-NMR spectrum of compound 13
Figure S23. ^1^H-NMR spectrum of compound 14

Figure S24. ^13^C-NMR spectrum of compound 14

Figure S25. ^1^H-NMR spectrum of compound 15

Figure S26. ^13^C-NMR spectrum of compound 15

Figure S27. ^1^H-NMR spectrum of compound 16

Figure S28. ^13^C-NMR spectrum of compound 16

Figure S29. ^1^H-NMR spectrum of compound 17

Figure S30. ^13^C-NMR spectrum of compound 17

Figure S1. HRMS spectrum of compound 3

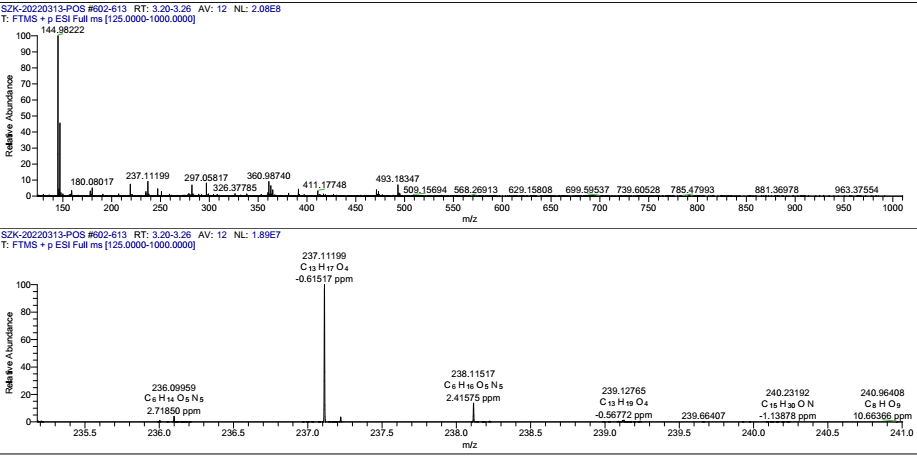

Figure S2. HRMS spectrum of compound 4

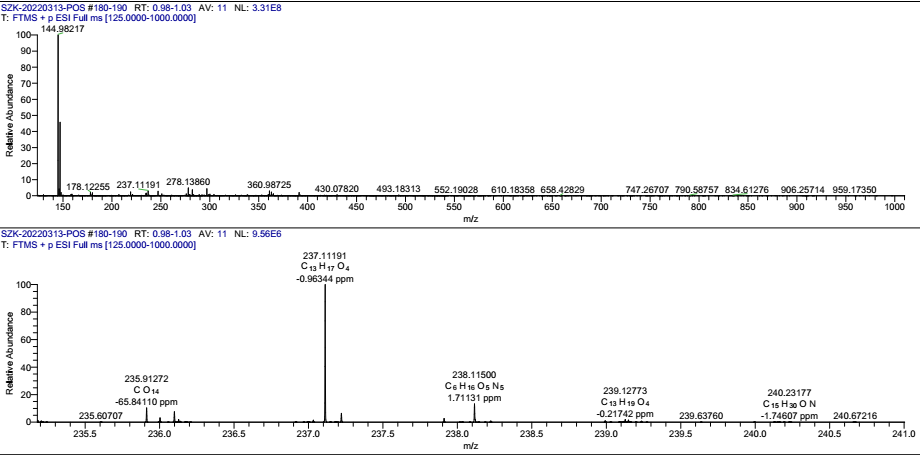

Figure S3. HRMS spectrum of compound 10

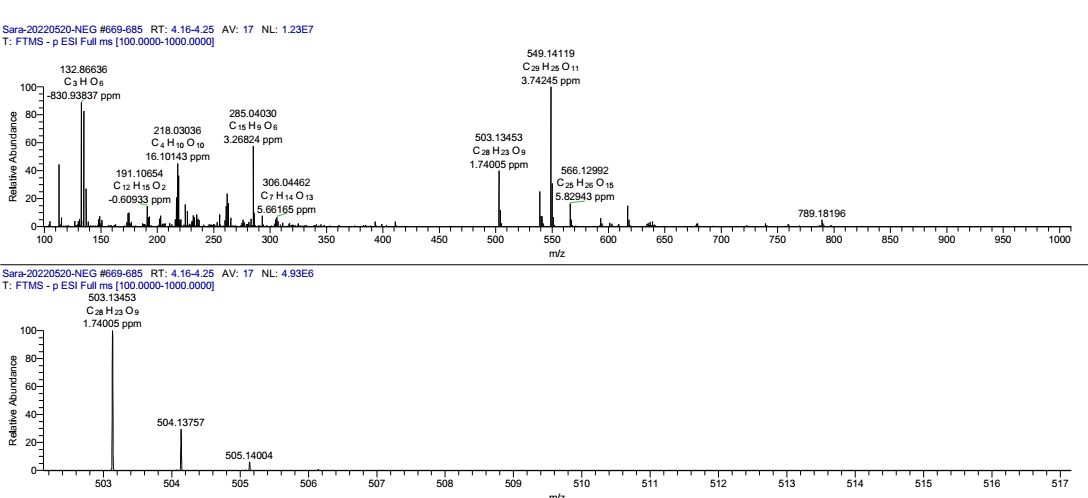

Figure S4. HRMS spectrum of compound 11

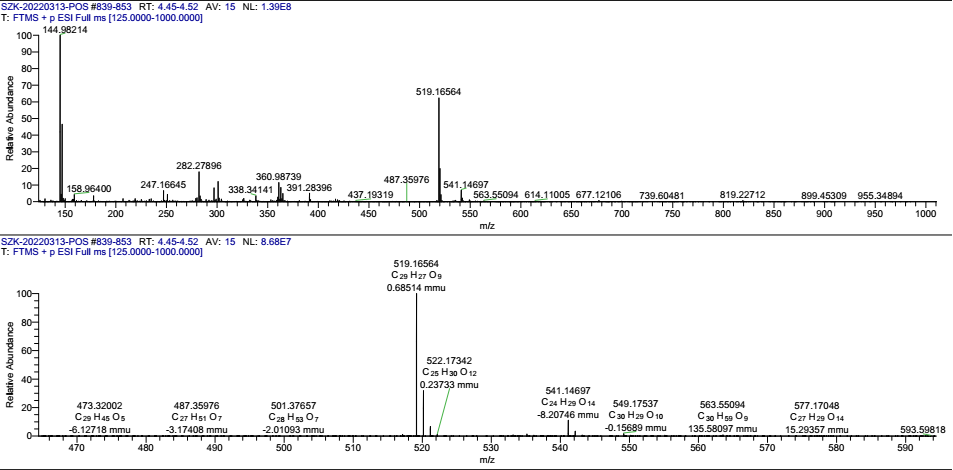

Figure S5. HRMS spectrum of compound 12

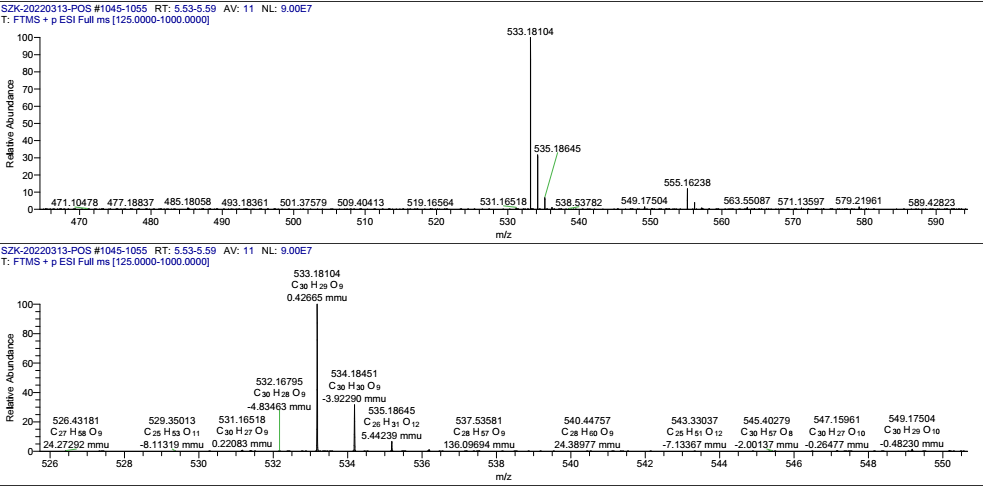

Figure S6. HRMS spectrum of compound 13

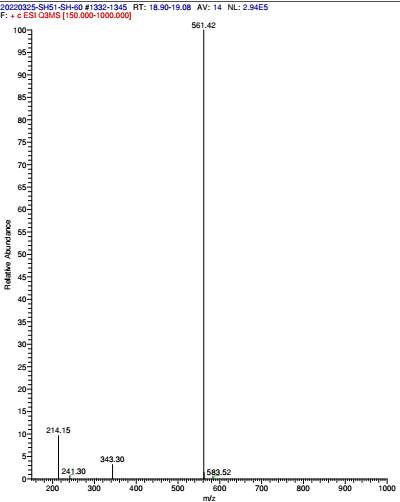

Figure S7. HRMS spectrum of compound 14

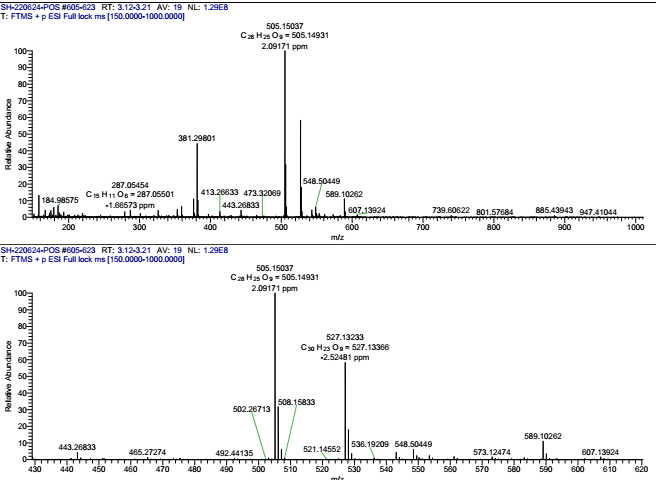

Figure S8. HRMS spectrum of compound 15

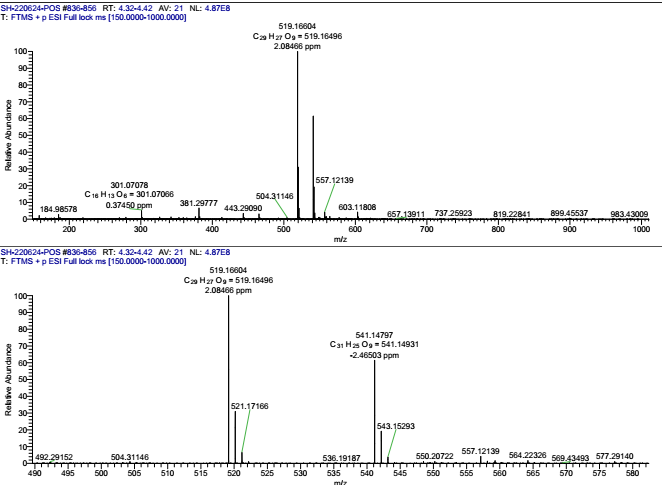

Figure S9. HRMS spectrum of compound 16

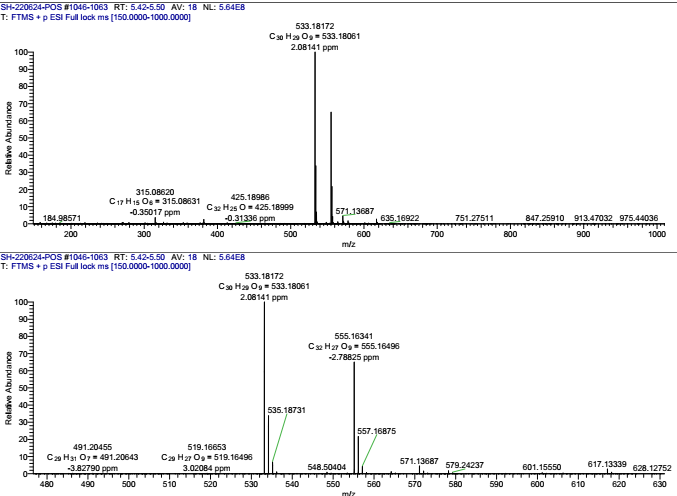

Figure S10. HRMS spectrum of compound 17

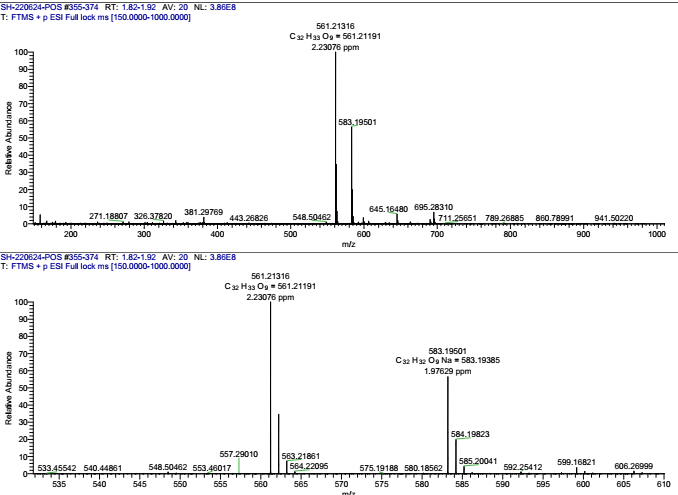

Figure S11. ^1^H-NMR spectrum of compound 3
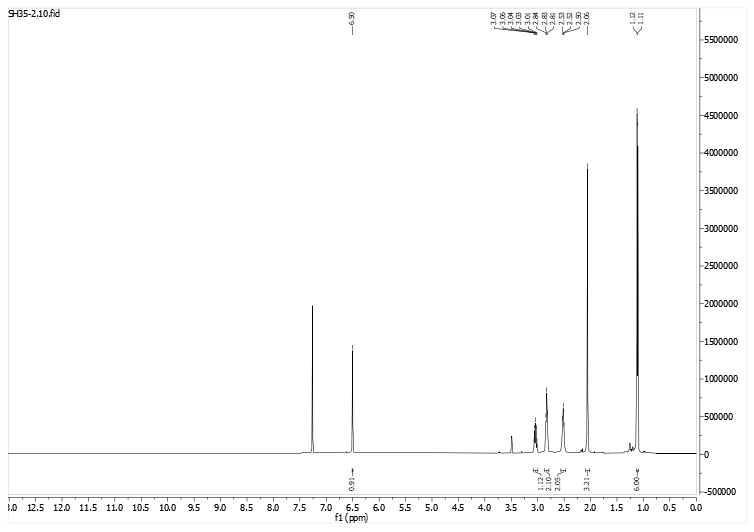

**Figure** S12. ^13^C-NMR spectrum of compound 3
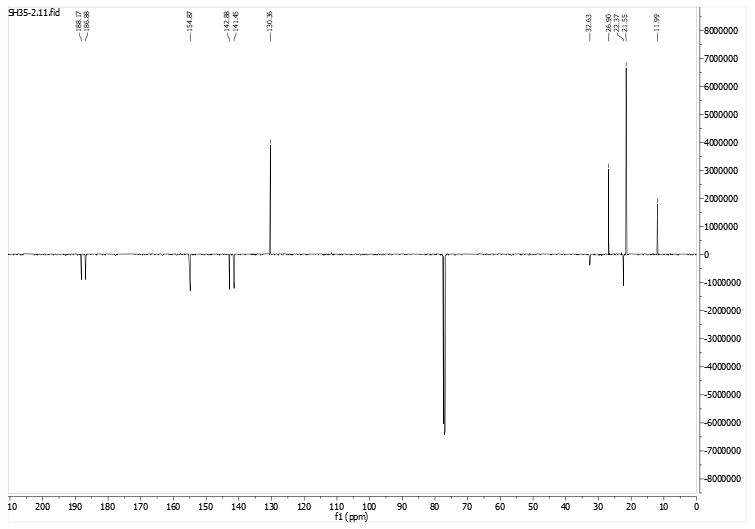

Figure S13. ^1^H-NMR spectrum of compound 4
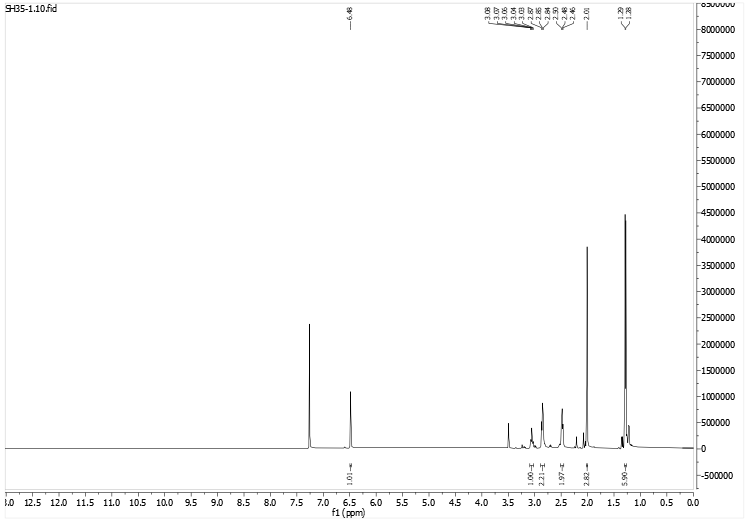

Figure S14. ^13^C-NMR spectrum of compound 4
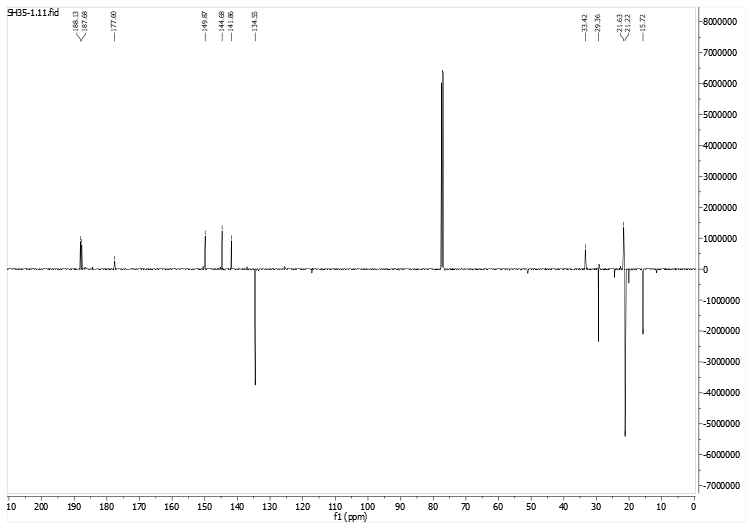

Figure S15. ^1^H-NMR spectrum of compound 10
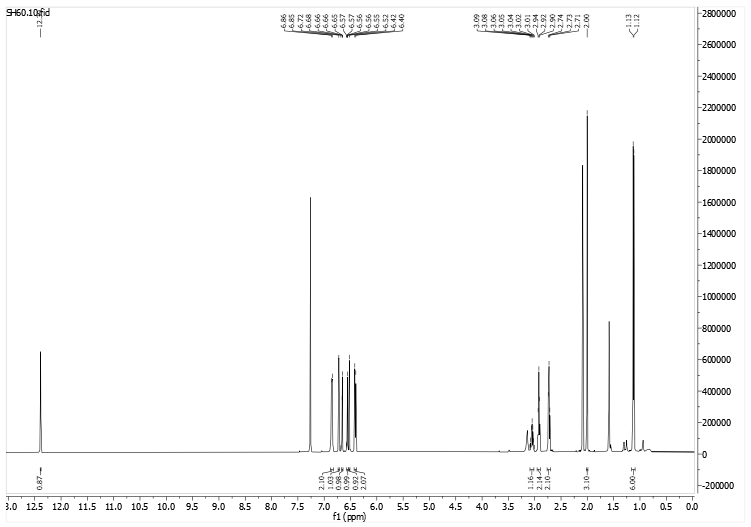

Figure S16. ^13^C-NMR spectrum of compound 10

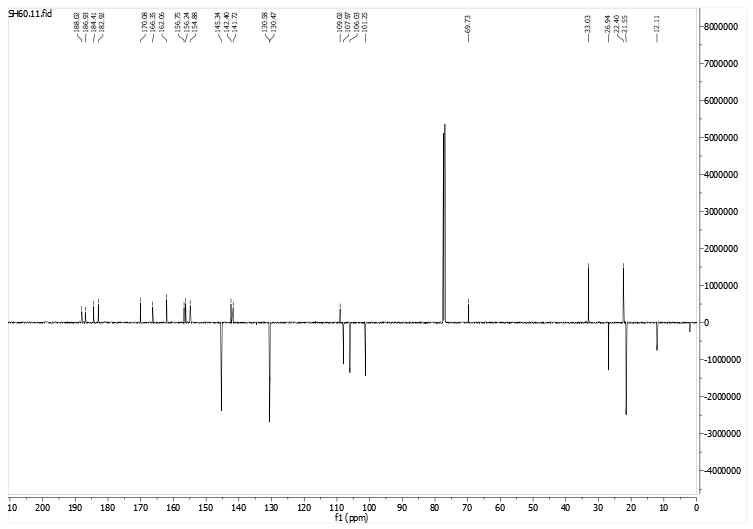

Figure S17. ^1^H-NMR spectrum of compound 11

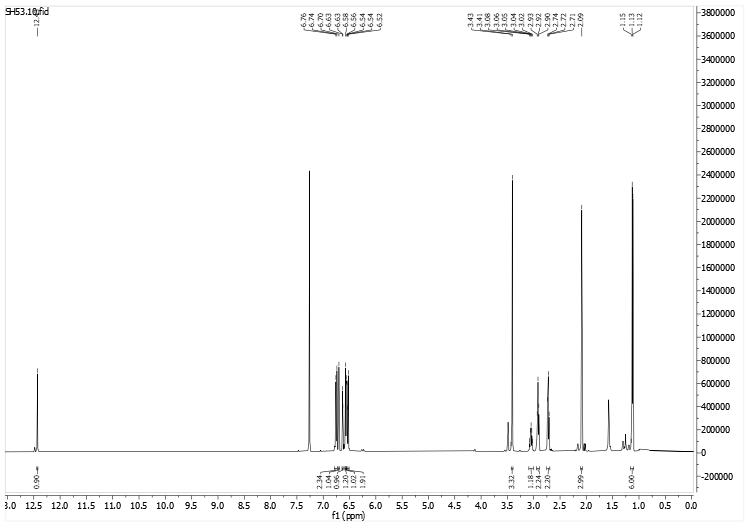

Figure S18. ^13^C-NMR spectrum of compound 11

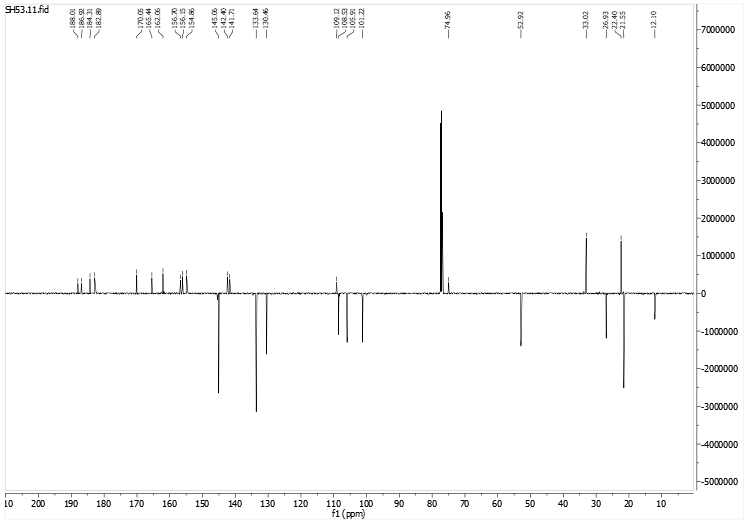

Figure S179 ^1^H-NMR spectrum of compound 12
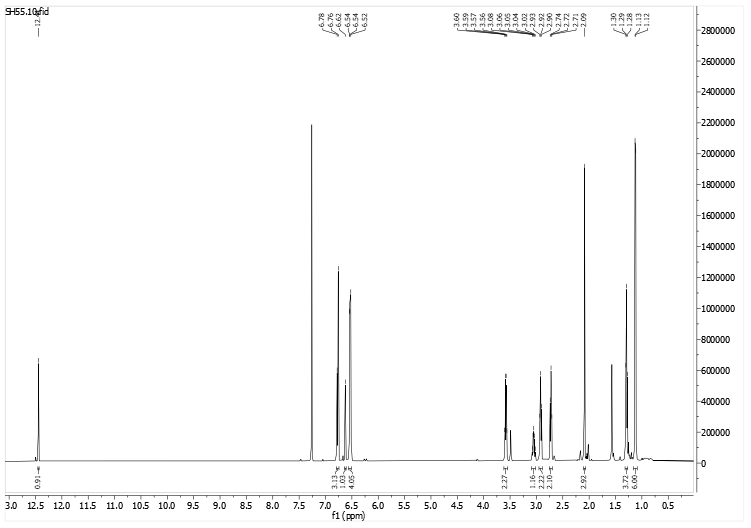

Figure S20 ^13^C-NMR spectrum of compound 12
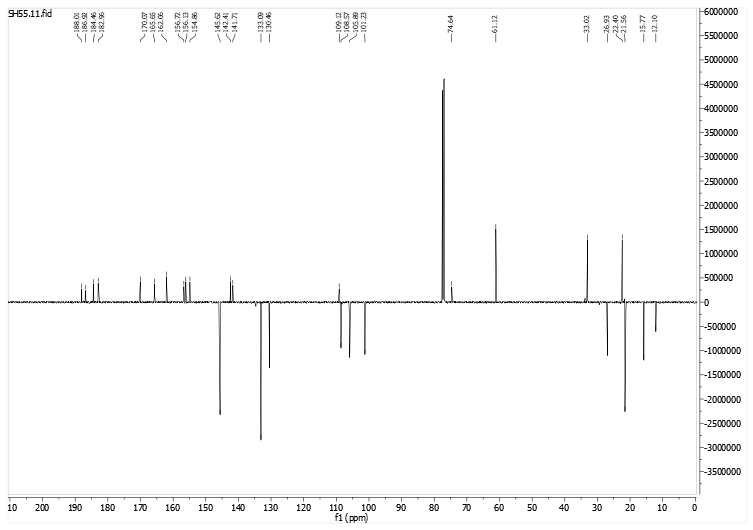

Figure S21. ^1^H-NMR spectrum of compound 13

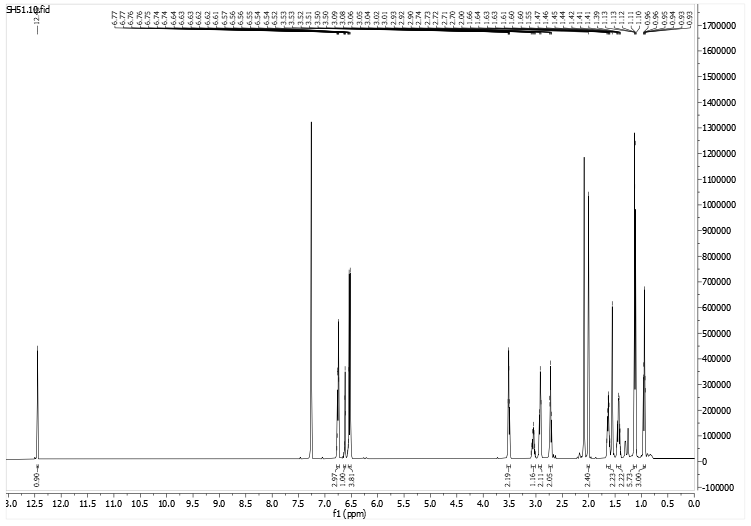

Figure S22. ^13^C-NMR spectrum of compound 13

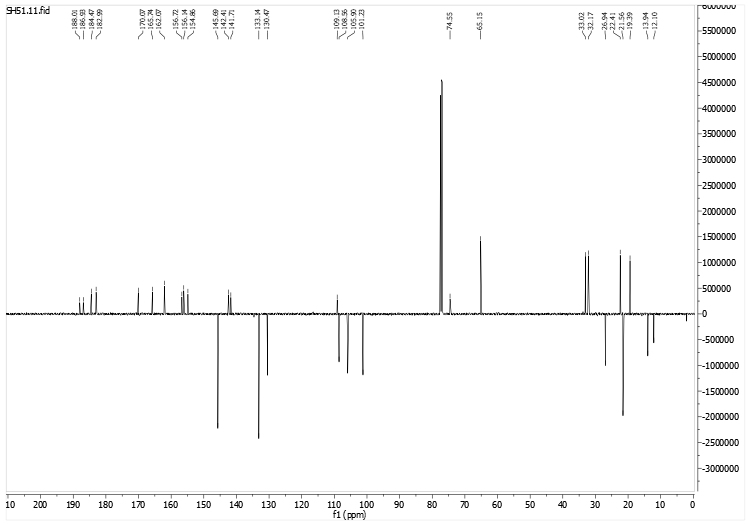

Figure S23. ^1^H-NMR spectrum of compound 14

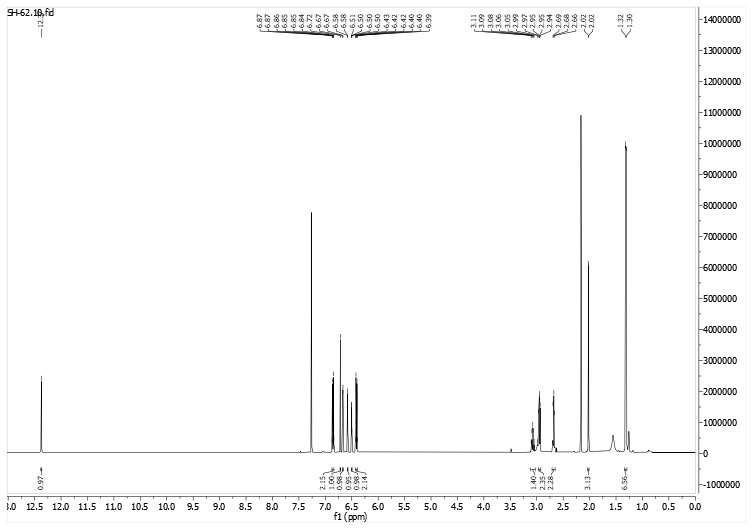

Figure S24. ^13^C-NMR spectrum of compound 14
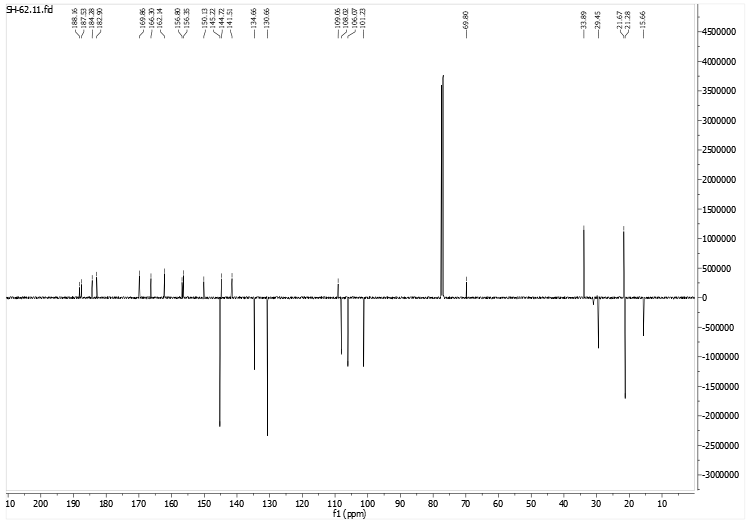

Figure S25. ^1^H-NMR spectrum of compound 15

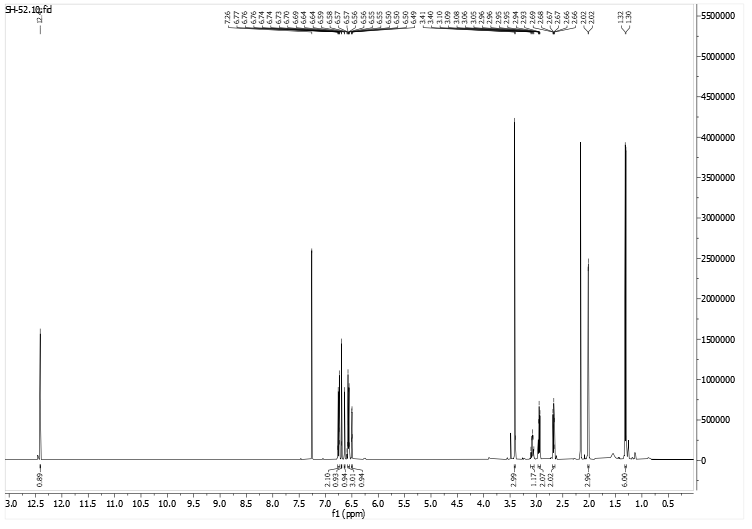

Figure S26. ^13^C-NMR spectrum of compound 15

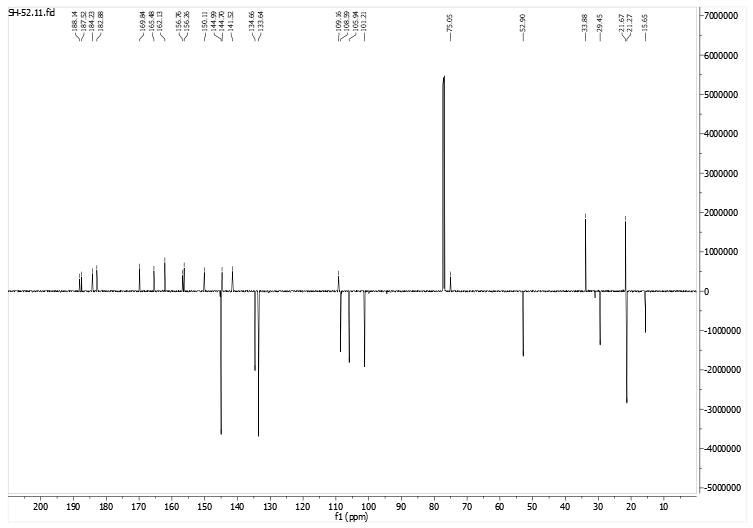

Figure S27. ^1^H-NMR spectrum of compound 16

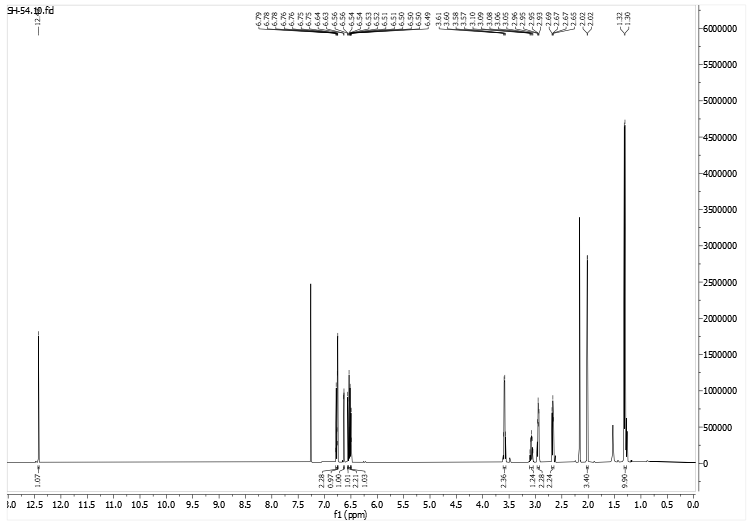

Figure S28. ^13^C-NMR spectrum of compound 16

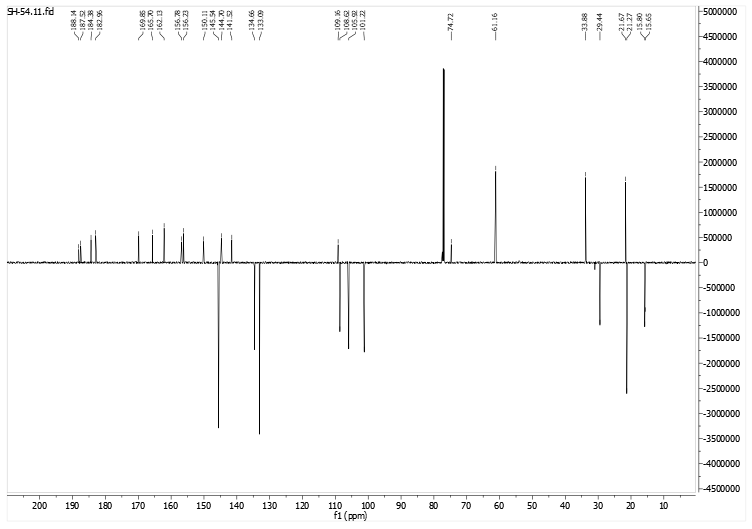

Figure S29. ^1^H-NMR spectrum of compound 17

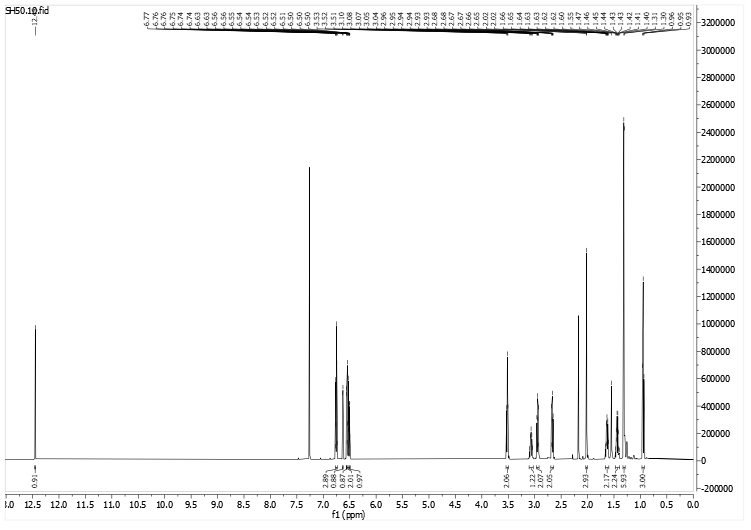

Figure S30. ^13^C-NMR spectrum of compound 17

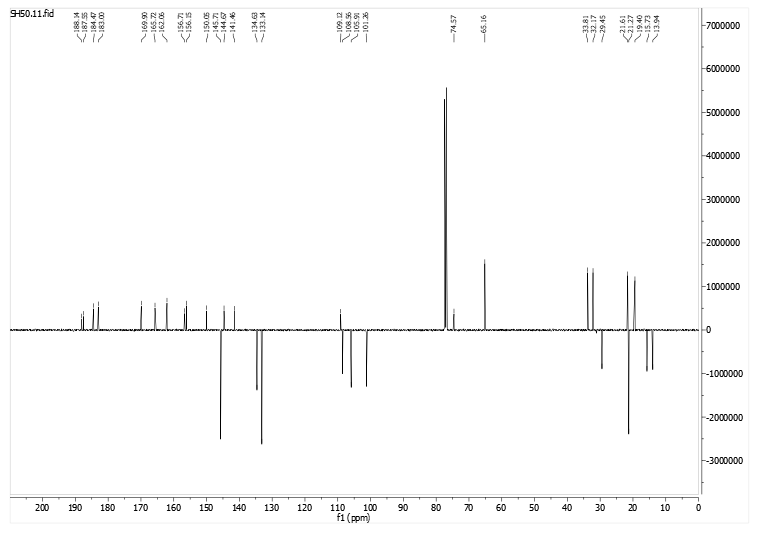
